# Engineered CRO-CD7 CAR-NK cells derived from pluripotent stem cells avoid fratricide and efficiently suppress human T-cell malignancies

**DOI:** 10.1101/2025.01.23.634617

**Authors:** Yunqing Lin, Ziyun Xiao, Fangxiao Hu, Xiujuan Zheng, Chenyuan Zhang, Yao Wang, Yanhong Liu, Dehao Huang, Zhiqian Wang, Chengxiang Xia, Qitong Weng, Leqiang Zhang, Yaoqin Zhao, Hanmeng Qi, Yiyuan Shen, Yi Chen, Fan Zhang, Jiaxin Wu, Pengcheng Liu, Jiacheng Xu, Lijuan Liu, Yanping Zhu, Jingliao Zhang, Wenbin Qian, Aibin Liang, Xiaofan Zhu, Tongjie Wang, Mengyun Zhang, Jinyong Wang

## Abstract

**Background:** T cell malignancies are highly aggressive hematological tumors with limited effective treatment options. CAR-NK cell therapy targeting CD7 has emerged as a promising approach for treating T-cell malignancies. However, conventional CAR-NK cell therapy faces the challenges of cell fratricide due to CD7 expression on both malignant cells and normal NK cells. Additionally, engineering CARs into human tissue-derived NK cells demonstrates heterogeneity, low transduction efficiency, and high manufacturing costs.

**Methods:** The human pluripotent stem cells (hPSCs) were genetically modified by knocking out the CD7 gene and introducing the CD7 CAR expression cassette to generate CD7 KO-CD7 CAR-hPSCs. These modified hPSCs were subsequently differentiated into CD7 KO-CD7 CAR-iNK cells using an efficient organoid induction method. The cytotoxicity of CD7 KO-CD7 CAR-iNK cells against CD7^+^ tumor cells was evaluated. Furthermore, we overexpressed the CXCR4 gene in CD7 KO-CD7 CAR-hPSCs and derived CXCR4-expressing CD7 KO-CD7 CAR-iNK (CRO-CD7 CAR-iNK) cells. The dynamics of CRO-CD7 CAR-iNK cells *in vivo* were tracked, and their therapeutic efficacy was assessed using human T-cell acute lymphoblastic leukemia (T-ALL) xenograft models.

**Results:** The CD7 KO-CD7 CAR-iNK cells derived from CD7 KO-CD7 CAR-hPSCs effectively avoided fratricide, demonstrated normal expansion, and exhibited potent and specific anti-tumor activity against CD7^+^ T-cell tumor cell lines and primary T-ALL cells. *CXCR4* overexpression in CRO-CD7 CAR-iNK cells improved their homing capacity and extended their persistence *in vivo*. The CRO-CD7 CAR iNK cells significantly suppressed tumor growth and prolonged the survival of T-ALL tumor-bearing mice.

**Conclusions:** Our study provides a reliable strategy for the large-scale generation of fratricide-resistant CD7 CAR-iNK cells with robust anti-tumor effects from hPSCs, offering a promising cell product to treat T-cell malignancy.

## Background

T cell malignancies are a type of highly aggressive hematological malignancy. Adult patients with relapsed or refractory T cell malignancies have poor disease outcomes with a 5-year overall survival rate lower than 20% [1–3]. Therefore, innovative therapies are needed for the treatment of T-cell malignancies. In recent years, chimeric antigen receptor (CAR)-T cell therapy targeting CD19 and BCMA has shown promising outcomes for relapsed or refractory B-cell malignancies and multiple myeloma [4, 5]. However, similar approaches to tackling T cell malignancies have been more challenging because of the shared expression of many targetable antigens between normal and malignant T cells, which results in fratricidal effects during CAR-T cell production.

CD7 is a transmembrane glycoprotein mainly expressed by T cells, natural killer (NK) cells, and their precursors [6, 7]. It also serves as a crucial surface marker for tumor cells in hematological malignancies and is expressed in 95% of lymphoblastic T-cell leukemias and lymphomas, as well as in a subset of peripheral T-cell lymphomas [8]. Therefore, immunotherapy that targets CD7 is capable of covering the majority of T-cell malignancy subtypes. Currently, multiple CD7 CAR-T cell therapies for T-cell leukemia and T-cell lymphoma are undergoing clinical trials [9]. Some studies have demonstrated the impressive short-term efficacy of allogeneic donor-derived anti-CD7 CAR T cells in an early-phase clinical trial involving patients with relapsed and/or refractory T-ALL [10, 11]. However, previous studies have reported that the expansion of CD7 CAR-T/CAR-NK cells was impaired due to fratricide, limiting their therapeutic potential [12]. To prevent fratricide in CAR-T and CAR-NK cells, strategies have been developed to remove surface CD7 antigen through genome editing or an intracellular protein expression blocker [12–17].

CAR-NK cells exhibit comparable effector functions to CAR-T cells and have emerged as a promising immunotherapy option due to their minimal toxicities and universality [18]. CD7 CAR-NK therapy is expected to provide a safer and more universally applicable immunotherapy for the treatment of T-cell malignancies. Nevertheless, tissue-derived NK cells face several challenges, including functional heterogeneities, low efficiencies, and high costs of gene engineering or editing. Human pluripotent stem cells (hPSCs) are promising cell sources to produce standardized, off-the-shelf NK cells, offering advantages such as unlimited cell sources and the ability to perform multiple genetic modifications. Pre-engineered CAR-hPSC-induced CAR-NK (CAR-iNK) cells have been reported to exhibit promising anti-tumor activity against both hematological and certain solid tumors [19–22].

Current treatment strategies for T cell malignancies have not demonstrated strong efficacy or long-term persistence of allogeneic CAR-NK cells after infusion [23]. Further efforts are required to enhance the therapeutic efficacy of CAR-NK therapy for T-cell malignancies. Studies have shown that *ex vivo* manipulation of NK cells results in the downregulation of CXCR4, which plays a crucial role in their homing to the bone marrow. Engineering strategies to stably equip effector NK cells with CXCR4 have shown promise in reinstating their ability to home to the bone marrow and enhancing their cytotoxic capacities [24–29].

In this study, we verified that umbilical cord blood (UCB)-derived CD7 CAR-NK cells attacked autologous NK cells and showed impaired expansion ability. To generate large-scale, fratricide-resistant CD7 CAR-iNK cells from hPSCs, we initially knocked out the CD7 gene at the hPSC stage (CD7 KO-hPSCs) using CRISPR/Cas12 technology and subsequently differentiated these CD7 KO-hPSCs into CD7 knockout NK cells (CD7 KO-iNK). The CD7 KO-iNK cells successfully avoided the fratricidal effect of CD7 CAR-NK cells. Then we overexpressed CD7 CAR in the CD7 KO-hPSCs and differentiated them into CD7 CAR-expressing CD7 KO-iNK (CD7 KO-CD7 CAR-iNK) cells. These cells exhibited restored expansion capacity and efficiently eliminated CD7^+^ T-cell tumor cell lines and primary T-ALL cells *in vitro*. To further enhance the *in vivo* persistence and tumor-killing potential of CD7 KO-CD7 CAR-iNK cells, CXCR4 was ectopically expressed in CD7 KO-CD7 CAR-hPSCs (CRO-CD7 CAR-hPSCs). CRO-CD7 CAR-iNK cells, derived from CRO-CD7 CAR-hPSCs, exhibited prolonged persistence *in vivo*. The CRO-CD7 CAR iNK cells significantly suppressed tumor growth and prolonged the survival of tumor-bearing mice. Therefore, we provided a reliable strategy to obtain large-scale generation of fratricide-resistant CRO-CD7 CAR iNK cells with robust anti-tumor effects from hPSCs, offering a promising cell product to treat T-cell malignancy.

## Methods

### Mice

B-NDG (NOD.CB17-*Prkdc^scid^Il2rg^tm1Bcgen^*/Bcgen) and B-NDG hIL15 (NOD.CB17-*Prkdc^scid^ Il2rg^tm1Bcgen^ Il15^tm1(IL15)Bcgen^*/Bcgen) mice were purchased from Biocytogen. Mice were housed in the SPF-grade animal facility of the Institute of Zoology, Chinese Academy of Sciences.

## Statistical analysis

All quantitative analyses were performed with SPSS (IBM SPSS Statistics 25). All data are represented as means ± SD, and the specific number (n) for each dataset is detailed in the figure legends. The two-tailed independent *t*-test, one-way ANOVA, two-way ANOVA, and Kruskal-Wallis tests were used to compare groups. Significance was defined as a P value of less than 0.05. Survival curves for tumor-bearing models were plotted using the Kaplan-Meier method and compared between groups using the logarithmic rank (Mantel-Cox) test. Statistical analysis was performed using the GraphPad Prism 9 software.

## Results

### Deletion of the CD7 gene enables iNK cells derived from hPSCs to avoid fratricide

To evaluate whether CD7 CAR expression in NK cells causes fratricide, we produced CD7 CAR-NK cells from UCB and assessed their cytotoxicity against NK cells **(Fig. 1A)**. CD3^-^ cells were isolated from the UCB mononuclear cells on day-6 and cocultured with K562-mIL21 cells for six days to expand NK cells. Flow cytometry analysis confirmed that 99.6% of the expanded NK cells expressed CD7 **(Additional file 1: Fig. S1A)**. On day 0, the expanded NK cells were transduced with CD7 CAR retrovirus (MOI = 5) using spin infection. The expression of CD7 CAR was assessed by flow cytometry two days after retroviral infection, revealing an infection rate of over 90% **(Fig. 1B)**. The CD7 CAR-NK cells were then cocultured with autologous uninfected NK cells derived from the same UCB sample to evaluate their fratricide effect **(Fig. 1C)**. The results demonstrated that CD7 CAR-NK cells effectively killed CD7^+^ NK cells, confirming fratricide **(Fig. 1D)**. Therefore, disrupting the surface expression of CD7 on NK cells to overcome fratricide was necessary for the production of clinical-scale CD7 CAR-NK cells. Previously, we developed an organoid induction method for generating NK (iNK) and CAR-iNK cells from hPSCs [22, 23, 30]. In this study, we aimed to generate CD7 CAR-iNK cells from hPSCs using this reliable scale-up method. Given the fratricide in CD7 CAR-iNK cells, we first knocked out the CD7 gene in hPSCs using CRISPR/Cas12 gene editing. Two gRNAs were designed to target sequences within the first intron and the second exon of the CD7 gene **(Fig. 1E)**. The gRNA- and CRISPR/Cas12i-encoding plasmids were electroporated into hPSCs. After electroporation, single-cell clones were picked and cultured for genomic verification of CD7 knockout (CD7-KO). Genome PCR confirmed the deletion of the CD7 gene in the hPSCs **(Additional file 1: Fig. S1B-D)**, indicating that we had successfully obtained CD7-KO hPSCs. CD7 KO-iNK cells were further generated from CD7 KO-hPSCs. The unmodified hPSCs were used as controls to monitor any disruptions in differentiation caused by genetic modification. Briefly, the lateral plate mesoderm cells with CD7 KO (CD7 KO-iLPM) were efficiently induced from CD7 KO-hPSCs after two days of monolayer induction. On day 2, the CD7 KO-iLPM and OP9 cells were mixed to prepare organoid aggregates. The organoids were plated into the transwell for 25-day NK cell induction **(Fig. 1F)**. More than 90% of resulting cells exhibited mature NK cell phenotypes (CD45^+^CD3^-^CD56^+^CD16^+/-^) on day 27. Over 96% of Ctrl-iNK cells expressed CD7, while CD7 KO-iNK cells showed no CD7 expression **(Fig. 1G)**. Flow cytometry analysis confirmed the absence of CD7 expression in CD7 KO-iNK cells. To determine whether CD7 KO-iNK cells could avoid fratricide effect, Ctrl-iNK and CD7 KO-iNK cells were cocultured with UCB-derived CD7 CAR-NK cells **(Fig. 1H)**. As expected, CD7 KO-iNK cells avoided CD7 CAR-NK cell-mediated cytotoxicity, whereas Ctrl-iNK cells, which expressed CD7, were significantly killed **(Fig. 1I)**. Therefore, deletion of the CD7 gene enables iNK cells to avoid attacks by CD7

**Fig. 1.**
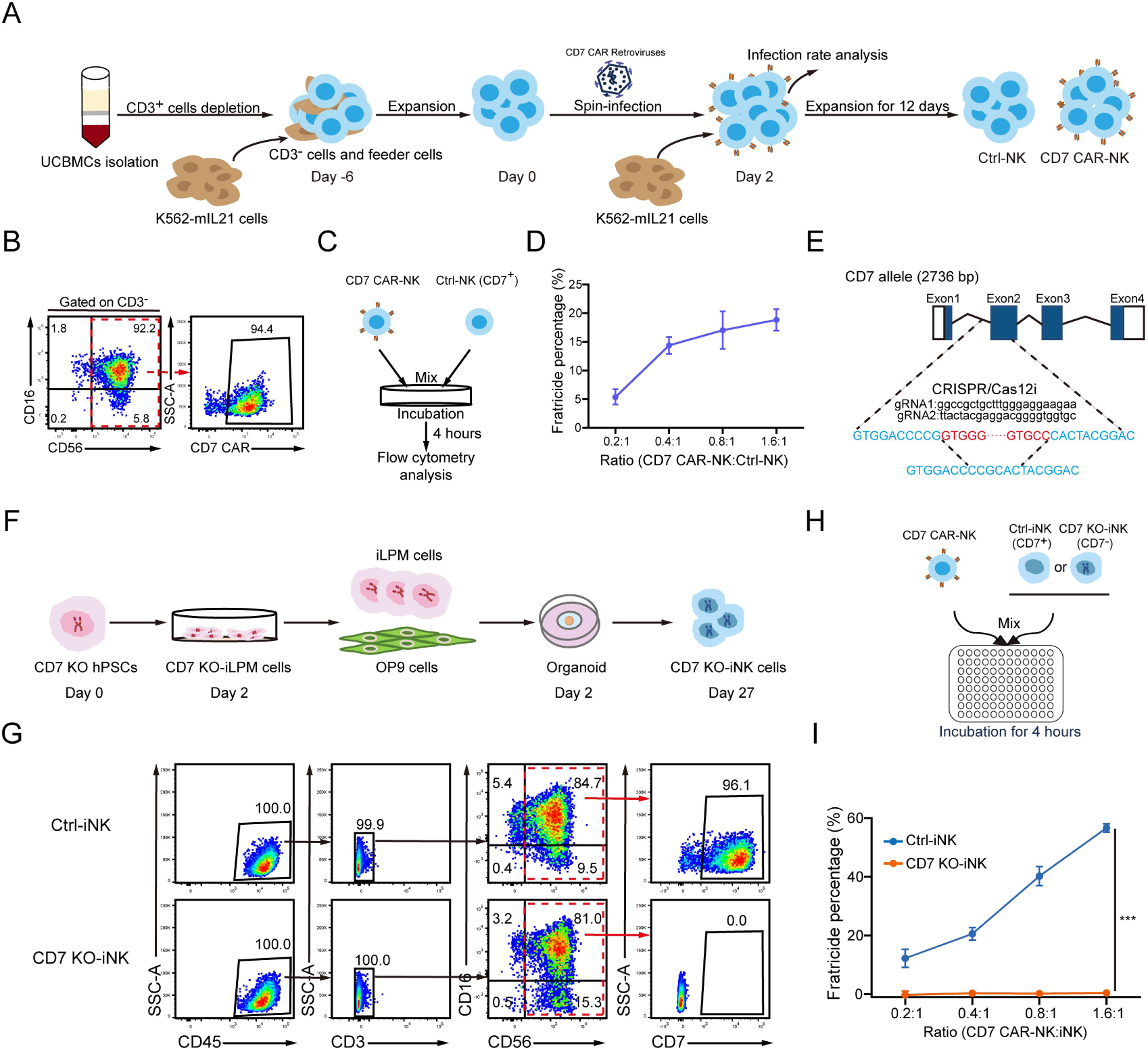
hPSC-derived iNK cells lacking CD7 expression avoid fratricide. **A** Schematic diagram showing the generation of CD7 CAR-NK cells from UCB. **B** Flow cytometry analysis showing the infection rate of CD7 CAR in CD3^-^CD56^+^ NK cells. The infection rate was analyzed 48 hours after transduction. **C** Schematic diagram of evaluating the cell lysis ratios of UCB-derived Ctrl-NK cells (uninfected with retrovirus) cocultured with CD7 CAR-NK cells. **D** Statistical analysis of the fratricide percentage of CD7 CAR-NK cells against Ctrl-NK cells at indicated E:T ratios (mean ± SD). n = 4 repeats. **E** Strategy for the knockout of CD7 gene in hPSCs. The exons of the CD7 gene are shown as blue boxes. Two gRNA sequences, gRNA1 and gRNA2, are shown in black. The deleted sequences are marked in red, and the sequences after editing are marked in blue. **F** Schematic diagram of CD7 KO-iNK cell induction. **G** Representative flow cytometry plots of CD7 KO-iNK cells (CD45^+^CD3^-^CD56^+^CD16^+/-^CD7^-^) on day 27. **H** Schematic diagram of evaluating the *in vitro* cytotoxic activity of CD7 CAR-NK cells against Ctrl-iNK cells or CD7 KO-iNK cells. **I** Statistical analysis of the fratricide percentage of CD7 CAR-NK cells against Ctrl-iNK cells or CD7 KO-iNK cells at indicated E:T ratios (two-tailed independent *t*-test, mean ± SD). n = 4 repeats. ***p < 0.001.

CAR-NK cells.

### Disruption of CD7 expression restores the expansion of CD7 CAR-iNK cells

Due to antigen-driven fratricide, UCB-derived CD7 CAR-NK cells exhibited poor expansion ability compared to the untransduced NK cells **(Fig. 2A)**. To achieve expandable and functional CD7 CAR-iNK cells from hPSCs, we engineered the CD7 CAR expression cassette into CD7 KO-hPSCs to construct the CD7 CAR-expressing CD7 KO-hPSCs (CD7 KO-CD7 CAR-hPSCs) **(Fig. 2B)**. The expression rates of CD7 CAR reached over 99% in CD7 KO-CD7 CAR-hPSCs after two rounds of sorting **(Additional file 1: Fig. S2A)**. CD7 KO-CD7 CAR-hPSCs were further differentiated into CD7 KO-CD7 CAR-iNK cells using the organoid induction method. Flow cytometry analysis showed that over 90% of CD7 KO-CD7 CAR-iNK cells expressed CD7 CAR while lacking CD7 expression **(Fig. 2C)**. The CD7 KO-CD7 CAR-iNK cells demonstrated comparable expansion folds to Ctrl-iNK cells, indicating that the deletion of CD7 successfully restored expansion capacity in CD7 CAR-iNK cells **(Fig. 2D)**. Furthermore, we assessed the expression of the crucial effector molecules in CD7 KO-CD7 CAR-iNK cells, including activating and inhibitory receptors (NKp30, NKp44, CD319, DNAM-1, CD96, NKG2A, and CD94), activating molecule (CD69), apoptosis-related ligands (TRAIL), and cytotoxic granules (GzmB and Perforin) **(Fig. 2E)**. The expression levels of these molecules were similar among Ctrl-iNK, CD7 KO-iNK, and CD7 KO-CD7 CAR-iNK cells. The CD7 KO-CD7 CAR-iNK cells were cocultured with UCB-NK, Ctrl-iNK, and CD7 KO-iNK cells to verify whether fratricide exists in the CD7 KO-CD7 CAR-iNK cells **(Fig. 2F)**. The results demonstrated that CD7 KO-CD7 CAR-iNK cells exerted strong cytotoxic activity against UCB-NK and Ctrl-iNK cells but exhibited no cytotoxicity against CD7 KO-iNK cells **(Fig. 2G)**. Taken together, we successfully generate expandable and fratricide-resistant CD7 KO-CD7 CAR-iNK cells from CD7 KO-CD7 CAR-hPSCs.

**Fig. 2.**
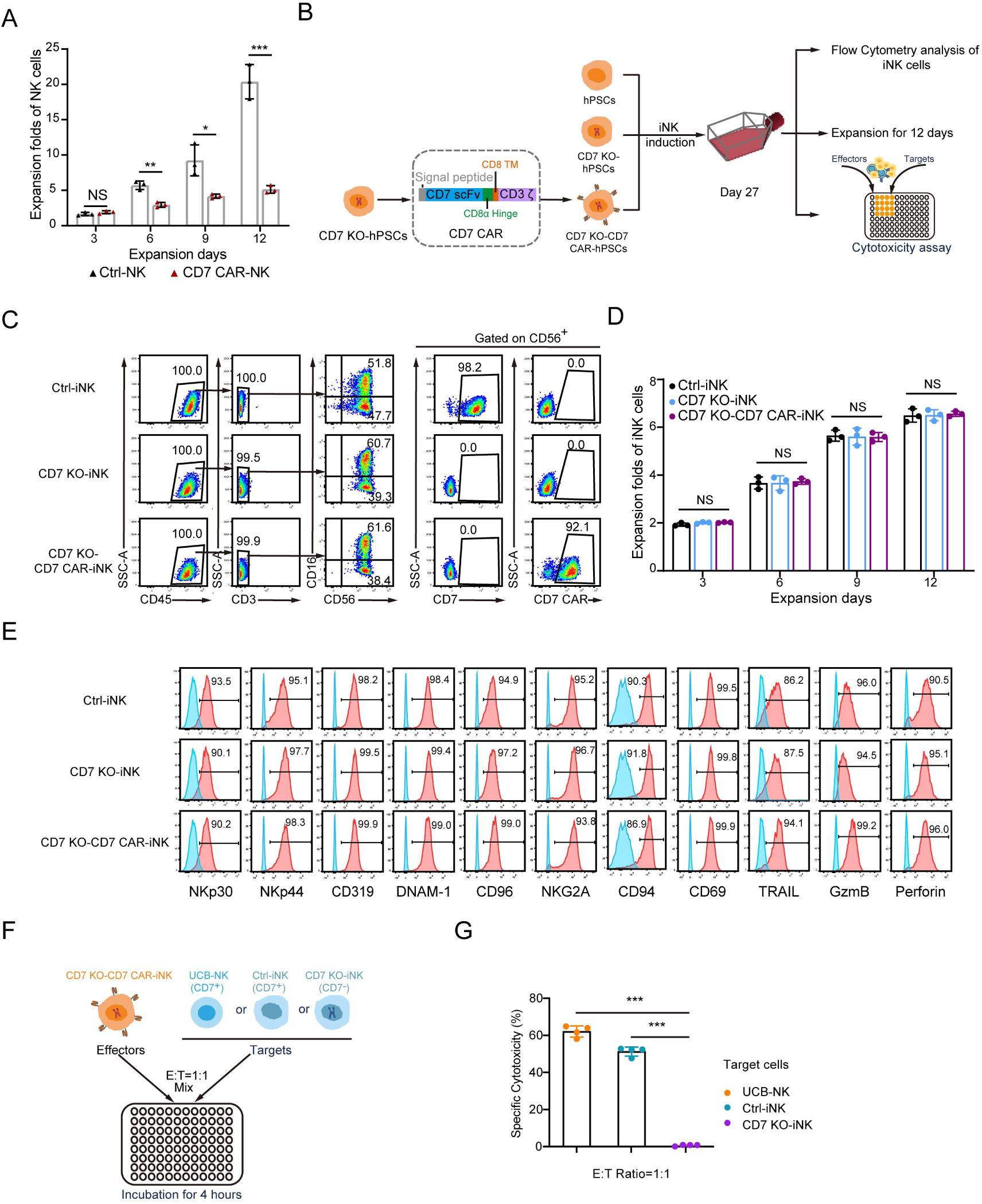
Disruption of CD7 expression restores the expansion of CD7 CAR-iNK cells. **A** Statistics analysis of the expansion folds of NK cells and CD7 CAR-NK cells derived from UCB on indicated expansion days (two-tailed independent *t*-test, mean ± SD). n = 3 repeats. NS, not significant (p > 0.05), *p < 0.05, **p < 0.01. ***p < 0.001. **B** Strategy diagram for the acquisition of CD7 KO-CD7 CAR-iNK cells and the detection of their expansion capacity and cytotoxic activity. **C** Representative flow cytometry plots of Ctrl-iNK, CD7 KO-iNK, and CD7 KO-CD7 CAR-iNK cells (CD45^+^CD3^-^CD56^+^CD16^+/-^CD7^-^CD7 CAR^+^) on day 27. **D** Statistics analysis of the expansion folds of Ctrl-iNK cells, CD7 KO-iNK cells, and CD7 KO-CD7 CAR-iNK cells on indicated expansion days (one-way ANOVA with Tukey’s multiple-comparison test). n = 3 repeats. NS, not significant (p > 0.05). **E** Flow cytometry histograms showing the expression levels of NKp30, NKp44, CD319, DNAM-1, CD96, NKG2A, CD94, CD69, TRAIL GzmB, and Perforin in Ctrl-iNK, CD7 KO-iNK, and CD7 KO-CD7 CAR-iNK cells. **F** Schematic diagram of evaluating the *in vitro* cytotoxic activity of CD7 KO-CD7 CAR-iNK cells against UCB-NK, Ctrl-iNK, and CD7 KO-iNK cells. **G** Statistical analysis of the cytotoxic activity of CD7 KO-CD7 CAR-iNK cells against UCB-NK, Ctrl-iNK, and CD7 KO-iNK cells at an E:T ratio of 1:1 (one-way ANOVA with Tamhane’s multiple-comparison test). n = 4 repeats. ***p < 0.001.

### CD7 KO-CD7 CAR-iNK cells show robust tumor-killing abilities *in vitro*

We subsequently performed tumor-killing assays to evaluate the cytotoxicity of CD7 KO-CD7 CAR-iNK cells against T-ALL cells. The Jurkat and CCRF-CEM cells, which both highly expressed CD7 protein **(Additional file 1: Fig. S2B)**, were used as targets for the specific cytotoxicity assay. Tumor cells (targets, T) were cocultured with either Ctrl-iNK cells or CD7 KO-CD7 CAR-iNK cells (effectors, E) at various E:T ratios **(Fig. 3A)**. As expected, CD7 KO-CD7 CAR-iNK cells efficiently targeted and induced apoptosis in Jurkat and CCRF-CEM cells within 4 hours, exhibiting significantly higher cytotoxicity compared to Ctrl-iNK cells **(Fig. 3B and C)**. To further assess the persistent cytotoxicity of CD7 KO-CD7 CAR-iNK cells in serial cytotoxic killing assays, we conducted three rounds of tumor killing assays with Jurkat or CCRF-CEM cells (E:T = 1:1) **(Fig. 3D)**. The results indicated that CD7 KO-CD7 CAR-iNK cells demonstrated a robust serial tumor-killing activity that was superior to that of Ctrl-iNK cells **(Fig. 3E and F)**. Then, NK cell stimulation assays were conducted by coculturing CD7 KO-CD7 CAR-iNK cells with Jurkat or CCRF-CEM cells at an E:T ratio of 1:1 for 4 hours. The expression of CD107a, a membrane protein associated with NK cell cytotoxic activity, and that of IFN-γ and TNF-α, NK cell cytotoxicity-related cytokines, was analyzed after incubation. As expected, the expression of CD107a in CD7 KO-CD7 CAR-iNK cells was significantly higher than that observed in Ctrl-iNK cells, indicating that CD7 KO-CD7 CAR-iNK cells released more cytotoxic granules **(Fig. 3G and H)**. Meanwhile, CD7 KO-CD7 CAR-iNK cells exhibited elevated production of IFN-γ and TNF-α compared to Ctrl-iNK cells when stimulated with tumor cells **(Fig. 3I-L)**. These results demonstrate the specific cytotoxic efficacy of CD7 KO-CD7 CAR-iNK cells.

**Fig. 3.**
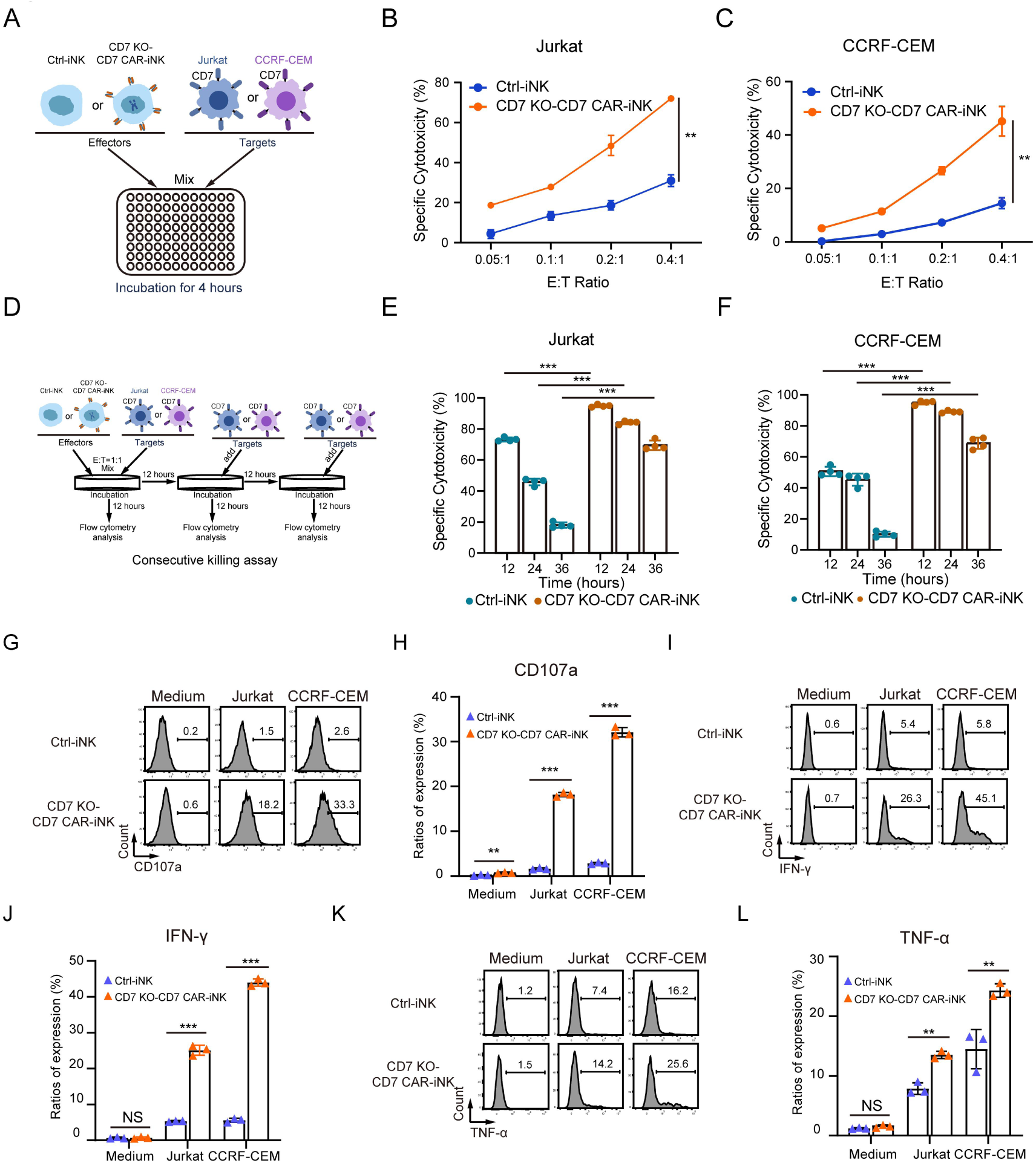
The cytotoxic activity of CD7 KO-CD7 CAR-iNK cells against CD7^+^ tumor cell lines *in vitro*. **A** Experimental design for the tumor-killing assay. The Ctrl-iNK and CD7 KO-CD7 CAR-iNK cells (effectors, E) were cocultured with Jurkat or CCRF-CEM tumor cells labeled with eFluor^TM^ 670 (targets, T), respectively. The E:T ratios include 0.05:1, 0.1:1, 0.2:1, and 0.4:1. **B-C** Cytotoxicity analysis of Ctrl-iNK cells and CD7 KO-CD7 CAR-iNK cells against Jurkat (**B**) or CCRF-CEM (**C**) tumor cells at the indicated E:T ratios after 4-hour incubation. Data are represented as means ± SD (n = 4). Kruskal-Wallis test was used for statistics. **p < 0.01. **D** Experimental design for the multiple rounds of tumor killing. The Ctrl-iNK and CD7 KO-CD7 CAR-iNK cells were respectively cocultured with Jurkat or CCRF-CEM tumor cells for 12 hours per round at the E:T ratio of 1:1. Fresh tumor cells were added to the Ctrl-iNK/CD7 KO-CD7 CAR-iNK cell residues incubated every other 12 hours. **E** Cytotoxicity analysis of three consecutive rounds of Jurkat cell killing by Ctrl-iNK or CD7 KO-CD7 CAR-iNK cells. Data are represented as means ± SD (n = 4). Two-tailed independent *t*-test was used for statistics. ***p < 0.001. **F** Cytotoxicity analysis of three consecutive rounds of CCRF-CEM cell killing by Ctrl-iNK or CD7 KO-CD7 CAR-iNK cells. Data are represented as means ± SD (n = 4). Two-tailed independent *t*-test was used for statistics. ***p < 0.001. **G-H** Assessment of CD107a expression by Ctrl-iNK or CD7 KO-CD7 CAR-iNK cells following 4 hours of coculture with Jurkat/CCRF-CEM tumor cells at the ratio of 1:1. Data are represented as means ± SD (n = 3). Two-tailed independent *t*-test was used for statistics. NS, not significant (p > 0.05), **p < 0.01, ***p < 0.001. **I-J** Measurement of IFN-γ production by Ctrl-iNK or CD7 KO-CD7 CAR-iNK cells in response to Jurkat/CCRF-CEM tumor cells. The Ctrl-iNK or CD7 KO-CD7 CAR-iNK cells were stimulated by Jurkat/CCRF-CEM at the ratio of 1:1 for 4 hours. Data are represented as means ± SD (n = 3). Two-tailed independent *t*-test was used for statistics. NS, not significant (p > 0.05), ***p < 0.001. **K-L** Evaluation of TNF-α production by Ctrl-iNK or CD7 KO-CD7 CAR-iNK cells in response to Jurkat/CCRF-CEM tumor cells. The Ctrl-iNK or CD7 KO-CD7 CAR-iNK cells were stimulated by Jurkat/CCRF-CEM at the ratio of 1:1 for 4 hours. Data are represented as means ± SD (n = 3). Two-tailed independent *t*-test was used for statistics. NS, not significant (p > 0.05), **p < 0.01.

### CD7 KO-CD7 CAR-iNK cells exhibit anti-tumor activity against CD7^+^ primary T-ALL cells

In addition to tumor cell lines, the cytotoxic potential of CD7 KO-CD7 CAR-iNK cells against primary T-ALL tumor cells was further evaluated. Freshly isolated mononuclear leukocytes from the bone marrow of three T-ALL patients were separately transplanted into B-NDG immunodeficient mice. Tumor progression was monitored weekly by analyzing the percentage of human CD45^+^CD7^+^ (huCD45^+^CD7^+^) tumor cells in PB using flow cytometry, starting four weeks after transplantation. Mice were sacrificed when the proportion of circulating huCD45^+^CD7^+^ cells exceeded 80% of the total PB mononuclear cell population. Subsequently, spleens from the B-NDG mice were harvested, processed into single-cell suspensions, and cocultured with iNK cells **(Fig. 4A)**. Six to nine weeks after transplantation, cells were obtained from the spleens of B-NDG immunodeficient mice in which huCD45^+^CD7^+^ cells in the PB reached over 80% **(Additional file 1: Fig. S3).** These cells were then stained to check CD7 expression. All samples showed nearly 100% of CD7^+^ cells **(Fig. 4B)**. These tumor cells obtained from the spleens were then cocultured with CD7 KO-CD7 CAR-iNK and Ctrl-iNK cells at defined E:T ratios. The results demonstrated that CD7 KO-CD7 CAR-iNK cells effectively targeted and eliminated primary T-ALL cells from all three patients **(Fig. 4C-E)**. Furthermore, the lysis of T-ALL tumor cells gradually increased over time, indicating the durable cytotoxicity of CD7 KO-CD7 CAR-iNK cells **(Fig. 4F-H)**. When the CD7 KO-CD7 CAR-iNK cells were cocultured with T-ALL cells at a 1:2 ratio, approximately 93.5%, 86.5%, and 64.5% of T-ALL cells from patient 1, patient 2, and patient 3 were eradicated within 12 hours, highlighting the potent effector function of the CD7 KO-CD7 CAR-iNK against primary tumors.

**Fig. 4.**
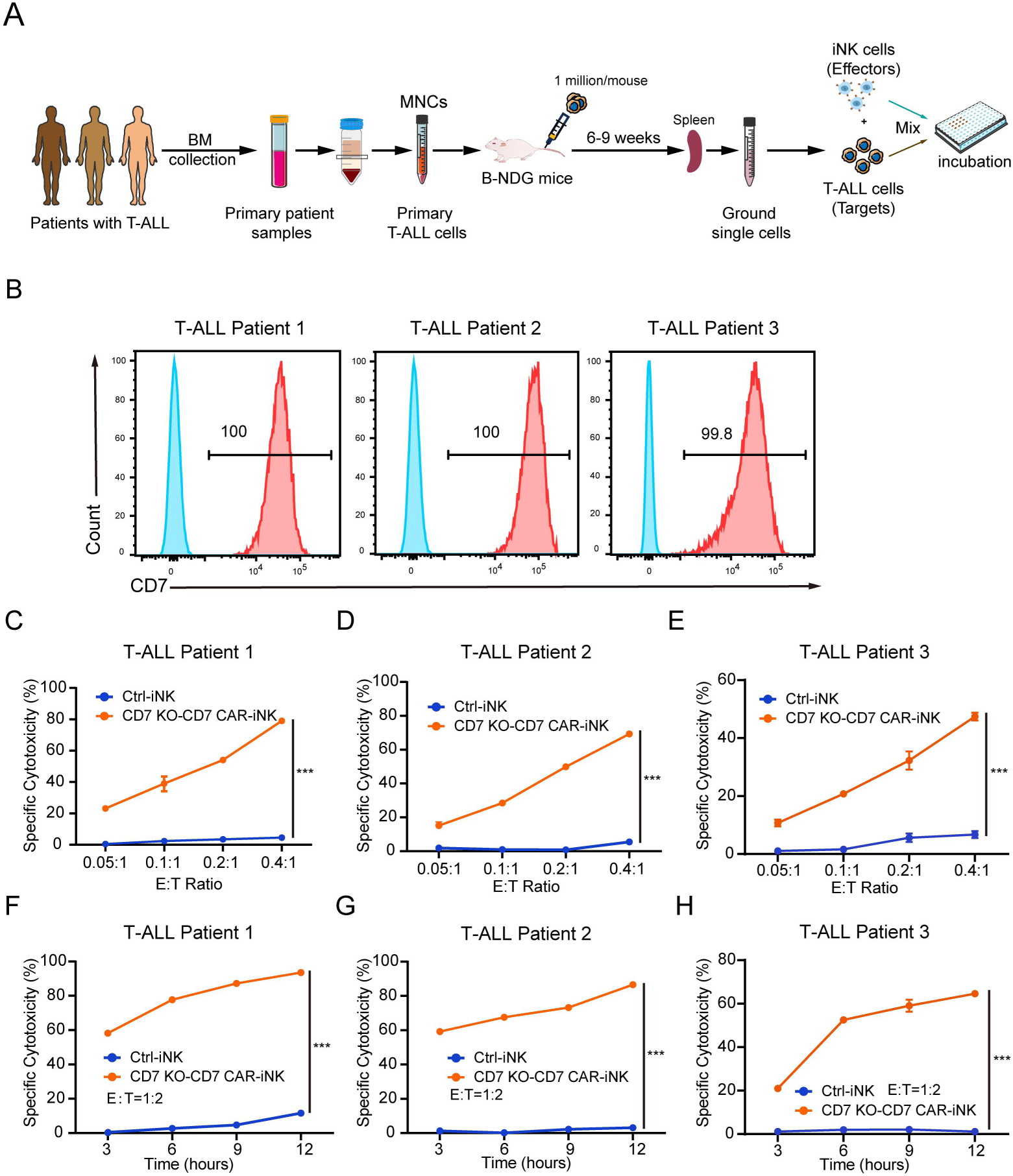
Anti-tumor activity of CD7 KO-CD7 CAR-iNK cells against primary T-ALL cells. **A** Experimental design for the T-ALL patient tumor-killing assay by hPSC-derived CD7 KO-CD7 CAR-iNK cells. Mononuclear cells were first isolated from the bone marrow of patients with T-ALL and transplanted into B-NDG immunodeficient mice (1×10^6^/mouse) via tail vein injection. Four weeks after transplantation, the proportion of primary tumor cells in the PB of B-NDG mice was monitored weekly using flow cytometry. Once the proportion of huCD45^+^CD7^+^ primary tumor cells exceeded 80% (approximately 6-9 weeks post-transplantation), the spleens of the B-NDG recipient mice were harvested and processed into single-cell suspensions. The resulting splenocytes were then cocultured with Ctrl-iNK or CD7 KO-CD7 CAR-iNK cells for tumor cytotoxicity assays. **B** Surface expression of CD7 in the splenic cells from B-NDG recipient mice measured by flow cytometry. **C-E** Cytotoxicity analysis of Ctrl-iNK or CD7 KO-CD7 CAR-iNK cells against T-ALL cells isolated from patient 1 (**C**), patient 2 (**D**), and patient 3 (**E**) at the indicated E:T ratios after 6-hour incubation. Data are represented as means ± SD (n = 4). Two-way ANOVA and Kruskal-Wallis tests were used for statistics. ***p < 0.001. **F-H** Cytotoxicity analysis of Ctrl-iNK or CD7 KO-CD7 CAR-iNK cells against T-ALL cells isolated from patient 1 (**F**), patient 2 (**G**), and patient 3 (**H**) at the E:T ratio of 1:2 after 3 h-, 6 h-, 9 h-, and 12 h- incubation. Data are represented as means ± SD (n = 4). Two-way ANOVA and Kruskal-Wallis tests were used for statistics. ***p < 0.001.

### CXCR4 improves the bone marrow homing capacity and prolongs the persistence of CD7 KO-CD7 CAR-iNK cells *in vivo*

To promote the migration of hPSC-derived CD7 KO-CD7 CAR-iNK cells to the bone marrow and further enhance their tumor-killing ability, we constructed CXCR4-overexpressing CD7 KO-CD7 CAR hPSCs (CRO-CD7 CAR-hPSCs). To enable real-time *in vivo* tracking, the CRO-CD7 CAR-hPSCs were engineered to express luciferase (CRO-CD7 CAR-luci-hPSCs). Luciferase-expressing CD7 KO-CD7 CAR hPSCs (CD7 KO-CD7 CAR-luci hPSCs) were also constructed as control **(Fig. 5A).** The CXCR4 and/or luciferase cassettes were integrated into the genomes of CD7 KO-CD7 CAR-hPSCs using the PiggyBac transposon system. The CXCR4 expression rates exceeded 98% in CRO-CD7 CAR hPSCs and CRO-CD7-CAR-luci hPSCs after two rounds of sorting **(Fig. 5B).** Bioluminescent imaging (BLI) verified the expression of luciferase in both CD7 KO-CD7 CAR-luci hPSCs and CRO-CD7 CAR-luci hPSCs **(Fig. 5C)**. CRO-CD7 CAR hPSCs, CD7 KO-CD7 CAR-luci hPSCs, and CRO-CD7 CAR-luci hPSCs were further differentiated into CRO-CD7 CAR-iNK, CD7 KO-CD7 CAR-luci-iNK, and CRO-CD7 CAR-luci-iNK cells using the organoid induction method. Flow cytometry analysis indicated the expression of CXCR4 in both CRO-CD7 CAR-iNK and CRO-CD7 CAR-luci-iNK cells **(Fig. 5D)**. To assess their *in vivo* distribution and persistence, CD7 KO-CD7 CAR-luci-iNK and CRO-CD7 CAR-luci-iNK cells were infused via tail vein into B-NDG hIL15 mice on day 0. BLI and flow cytometry analysis were performed at designated time points to monitor the dynamic distribution and persistence of infused cells **(Fig. 5E)**. BLI at the different time points post-infusion revealed that both cell fractions were initially trapped predominantly in the lungs; however, after 24 hours, they assumed distinctly different anatomical localizations. The CRO-CD7 CAR-luci-iNK cells were found mainly in bone marrow niches including the sternum, vertebrae, and bones of the lower extremities after 24 hours, and they could be maintained *in vivo* for more than 60 days **(Fig. 5F and G)**. Flow cytometry analysis of the PB from iNK-infused B-NDG hIL15 mice showed that the proportion of infused huCD45^+^ cells increased from about 0.6% at 7 days after transplantation to 11% - 15% at 14 days after transplantation and then gradually decreased **(Fig. 5H)**. Notably, the persistence of CRO-CD7 CAR-luci-iNK cells was significantly prolonged compared to the CD7 KO-CD7 CAR-luci-iNK group, which persisted for less than 28 days **(Fig. 5F-H)**. Taken together, these data indicate that CXCR4 overexpression enhances the homing capacity of CD7 KO-CD7 CAR-iNK cells to bone marrow compartments following adoptive transfer and extends their persistence *in vivo*.

**Fig. 5.**
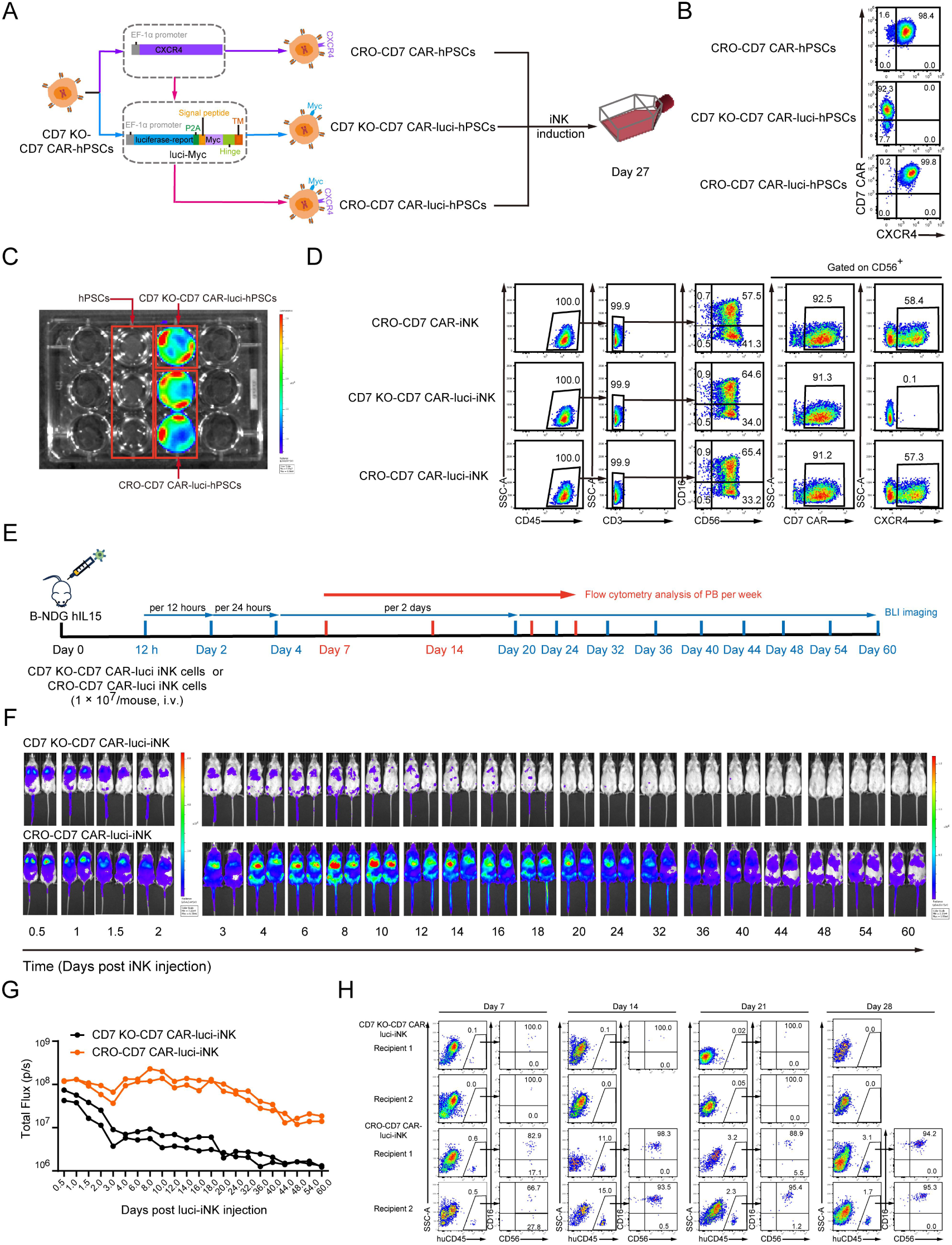
Dynamic analysis of CRO-CD7 CAR-iNK cells in the B-NDG hIL15 mice after infusion. **A** Strategy for the construction of CXCR4-CD7 KO-CD7 CAR-hPSCs (CRO-CD7 CAR-hPSCs), CD7 KO-CD7 CAR-luciferase-hPSCs (CD7 KO-CD7 CAR-luci-hPSCs), and CXCR4-CD7 KO-CD7 CAR-luciferase-hPSCs (CRO-CD7 CAR-luci-hPSCs). **B** Flow cytometry analysis of CXCR4 and CD7 CAR expression in CRO-CD7 CAR-hPSCs, CD7 KO-CD7 CAR-luci-hPSCs, and CRO-CD7 CAR-luci-hPSCs. **C** BLI imaging of the CD7 KO-CD7 CAR-luci-hPSCs and CRO-CD7 CAR-luci-hPSCs. **D** Representative flow cytometry plots of CRO-CD7 CAR-iNK, CD7 KO-CD7 CAR-luci-iNK, and CRO-CD7 CAR-luci-iNK cells on day 27. **E** Schematic diagram of *in vivo* studies with CD7 KO-CD7 CAR-luci-iNK and CRO-CD7 CAR-luci-iNK cells in B-NDG hIL15 mice. The CD7 KO-CD7 CAR-luci-iNK and CRO-CD7 CAR-luci-iNK cells were injected into B-NDG hIL15 mice (1 × 10^7^ cells/mouse) via the tail vein on day 0, respectively. BLI was performed at indicated time points. **F** BLI images showing the presence of CD7 KO-CD7 CAR-luci iNK or CRO-CD7 CAR-luci iNK cells in B-NDG hIL15 mice (n = 2 mice in each group) at the indicated time points. **G** Total flux (p/s) of the CD7 KO-CD7 CAR-luci-iNK or CRO-CD7 CAR-luci-iNK cells injected B-NDG hIL15 mice measured at the indicated time points (n = 2 mice in each group). **H** Flow cytometry analysis of CD7 KO-CD7 CAR-luci-iNK or CRO-CD7 CAR-luci-iNK cells in the PB of B-NDG hIL15 mice at the indicated time points.

### CRO-CD7 CAR-iNK cells suppress the tumor progress in xenograft animals

To further assess the therapeutic effects of the CRO-CD7 CAR-iNK cells on tumor cells *in vivo*, we established the T-ALL xenograft animal models by transplanting the luciferase-expressing CCRF-CEM (CCRF-CEM-luci) cells into B-NDG hIL15 immune-deficient mice. The B-NDG hIL15 mice were injected with CCRF-CEM-luci tumor cells (1 × 10^5^ cells/mouse) via tail vein on day 0 and received 1.0 Gy irradiation on day 1. Subsequently, Ctrl-iNK or CRO-CD7 CAR-iNK cells were intravenously injected into the tumor-bearing animals (1 × 10^7^ cells/mouse) on day 1, day 4, and day 7, while phosphate-buffered saline (PBS) was intravenously injected as control. BLI was performed weekly to capture the kinetics of tumor growth **(Fig. 6A)**. The data showed that tumor burdens of the PBS group and Ctrl-iNK group became increasingly severe, as indicated by the radiance and the value of total flux **(Fig. 6B and C)**. Eventually, both the PBS and Ctrl-iNK groups of mice needed ethical euthanasia due to the heavy tumor burden. However, the CRO-CD7 CAR-iNK cells showed stronger tumor-killing ability in xenograft animals with lower radiance and total flux measurements **(Fig. 6B and C)**. The CRO-CD7 CAR-iNK cell-treated mice survived significantly longer than the Ctrl-iNK cell-treated mice and the PBS-treated mice (Tumor + PBS: 29 days; Tumor + Ctrl-iNK: 30 days; Tumor + CRO-CD7 CAR-iNK: 36 days; p < 0.01) **(Fig. 6D)**. In conclusion, these results show that the CRO-CD7 CAR-iNK cells can efficiently suppress tumor development and prolong the survival of tumor-bearing mice, highlighting their potential as a therapeutic option for T-ALL treatment.

**Fig. 6.**
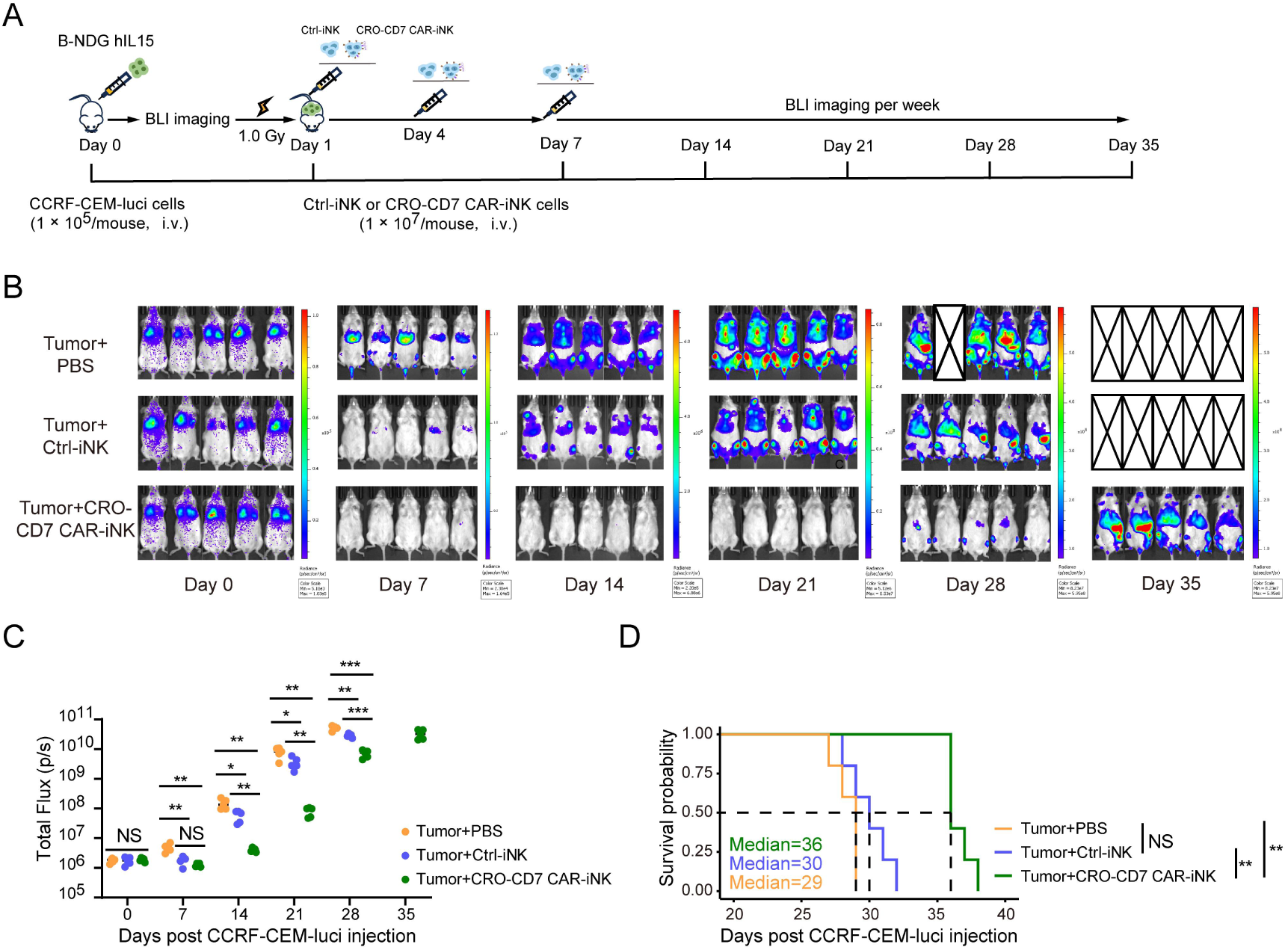
Suppression of human T-cell leukemia progress in xenograft models by CRO-CD7 CAR-iNK cells. **A** Schematic diagram of *in vivo* studies with luciferase-expressing CCRF-CEM cells (CCRF-CEM-luci) in mouse xenograft models. The CCRF-CEM-luci tumor cells were injected into B-NDG hIL15 mice (1 × 10^5^/mouse) via the tail vein on day 0. The mice were irradiated (1.0 Gy) on day 1. The equivalent of 1 × 10^7^ Ctrl-iNK or CRO CD7 CAR-iNK cells were infused into each animal on day 1, day 4, and day 7. Bioluminescent image (BLI) was performed every week. **B** BLI images of the xenograft models at the indicated time points (Tumor+PBS, Ctrl-iNK+PBS, and CRO-CD7 CAR-iNK+PBS, n = 5 mice in each group). The radiance indicates tumor burden. **C** Total flux (p/s) of the xenograft models measured at the indicated time points (n = 5 mice in each group). Two-tailed independent *t*-test was used for statistics. NS, not significant (p > 0.05), *p < 0.05, **p < 0.01. ***p < 0.001. **D** Kaplan-Meier survival curves of the xenograft models (n = 5 mice in each group). Median survival: Tumor + PBS, 29 days; Tumor + Ctrl-iNK, 30 days; Tumor + CRO-CD7 CAR-iNK, 36 days. The logarithmic rank (Mantel-Cox) test was used for statistics. NS, not significant, **p < 0.01.

## Discussion

In this study, we developed hPSC-derived CRO-CD7 CAR-iNK cells that avoid fratricide and show improved persistence *in vivo*. CD7 CAR-NK cells derived from UCB exhibited impaired expansion due to fratricide. By knocking out the CD7 gene and ectopically expressing CD7 CAR in hPSCs, we successfully generated expandable and functional CD7 KO-CD7 CAR-iNK cells using an organoid culture method. These CD7 KO-CD7 CAR-iNK cells effectively avoided CD7-antigen-induced fratricide and exhibited robust cytotoxic activity against CD7^+^ tumor cells *in vitro*, particularly primary tumor cells derived from T-ALL patients. Furthermore, CRO-CD7 CAR-iNK cells were generated from CXCR4-overexpressing CD7 KO-CD7 CAR-hPSCs to improve therapeutic efficacy. CXCR4 overexpression improved the bone marrow homing capacity and significantly prolonged the persistence of CRO-CD7 CAR-iNK cells *in vivo*. CRO-CD7 CAR-iNK effectively inhibited tumor progression in xenograft mice and significantly prolonged their survival.

CD7 is expressed not only on T cell malignancies and acute myeloid leukemia but also on most T cells and NK cells. Consequently, CAR-NK cells targeting CD7 would induce severe fratricide among themselves, leading to reduced proliferation and diminished therapeutic potential. Previous studies have proposed that one approach to overcome fratricide is to delete the CD7 gene from T cells [31]. The absence of CD7 in CAR-T cells does not impair their functionality. Therefore, we adopted the CD7 knockout strategy to prevent fratricide in CD7 CAR-NK cells. Instead of performing gene editing at the mature T or NK cell stages [15, 32], we conducted the gene editing at the hPSC stage. NK cells generated from gene-modified hPSCs exhibited the advantages of homogeneity, higher gene editing efficiency, and lower cost of CAR engineering. The CD7 KO-CD7 CAR-iNK cells exhibited comparable expansion ability and robust cytotoxic activity, indicating that the knockout of CD7 did not affect the development or function of hPSC-derived iNK cells.

CD7 CAR-T cells can be used to treat CD7-positive leukemias, including T-ALL, T-cell lymphoblastic lymphoma, and certain subtypes of acute myeloid leukemia [10, 14, 33, 34]. The CRO-CD7 CAR-iNK cells are capable of avoiding fratricide and demonstrating robust cytotoxic activity against CD7^+^ tumor cells. These features make them highly promising for the treatment of the above-mentioned hematological malignancies. Recent research has also demonstrated that iPSC-derived CD70 CAR-NK cells, which selectively target activated CD70^+^ T cells, offer a novel strategy for preventing allogeneic T cell rejection [35]. Similarly, CRO-CD7 CAR-iNK cells can target and deplete recipient CD7^+^ T cells, thereby enhancing their *in vivo* persistence and therapeutic efficacy. Moreover, the infusion of CRO-CD7 CAR-iNK cells has potential applications in the treatment of T cell-mediated autoimmune diseases, such as type 1 diabetes and multiple sclerosis [36, 37], by selectively eliminating pathogenic T cells.

Despite significant progress in treating T-ALL, long-term suppression of tumor progression remains a challenge, with few drugs or treatments achieving durable efficacy [38]. CXCR4 can significantly enhance the *in vivo* distribution and improve the persistence of iNK cells, thus enhancing the anti-tumor ability [39]. We have previously reported that CXCR4-expressing NK progenitor cells derived from hPSCs can effectively eliminate minimal residual disease in tumor models [29]. Therefore, combining CD7 CAR-iNK progenitor cells with chemotherapeutic drugs holds promise for effectively eliminating minimal residual disease in T-ALL models.

## Conclusions

In conclusion, CRO-CD7 CAR-iNK cells derived from hPSCs avoid fratricide and exhibit improved persistence and robust anti-tumor abilities *in vivo*. This study offers insights into the clinical potential of hPSC-derived CD7 CAR-iNK cells for the treatment of T cell malignancies.

## List of abbreviations

hPSCs: Human pluripotent stem cells
T-ALL: T-cell acute lymphoblastic leukemia
UCB: Umbilical cord blood
CAR: Chimeric antigen receptor
CD7 KO-hPSC: CD7 knockout hPSCs
CD7 KO-iNK: CD7 knockout iNK
CD7 KO-CD7 CAR-hPSCs: CD7 CAR-expressing CD7 KO-hPSCs
CD7 KO-CD7 CAR-iNK: CD7 CAR-expressing CD7 KO-iNK
CRO-CD7 CAR-hPSCs: CXCR4-expressing CD7 KO-CD7 CAR-hPSCs
CRO-CD7 CAR-iNK: CXCR4-expressing CD7 KO-CD7 CAR-iNK
CD7 KO-CD7 CAR-luci-hPSCs: CD7 KO-CD7 CAR-luciferase-hPSCs
CRO-CD7 CAR-luci-hPSCs: CXCR4-CD7 KO-CD7 CAR-luciferase-hPSCs
CD7 KO-CD7 CAR-luci-iNK: CD7 KO-CD7 CAR-luciferase-iNK
CRO-CD7 CAR-luci-iNK: CXCR4-CD7 KO-CD7 CAR-luciferase-iNK
B-NDG: NOD.CB17-*Prkdc^scid^Il2rg^tm1Bcgen^*/Bcgen
B-NDG hIL15: NOD.CB17-*Prkdc^scid^ Il2rg^tm1Bcgen^ Il15^tm1(IL15)Bcgen^*/Bcgen
scFv: Single-chain variable fragment
MNCs: Mononuclear cells
E:T ratio: Effector:Target ratio
IFN-γ: Interferon gamma
TNF-α: Tumor necrosis factor α
NK: Natural killer
PB: Peripheral blood
BLI: Bioluminescence imaging
PBS: Phosphate-buffered saline.

## Declarations

## Ethics approval

Experiments and handling of mice were conducted under the Institutional Animal Care and Use Committee of the Institute of Zoology, Chinese Academy of Sciences. The studies involving humans were approved by the Biomedical Research Ethics Committee of the Institute of Zoology, Chinese Academy of Sciences. The use of patient samples was conducted in accordance with the provisions of the Declaration of Helsinki. All patient samples were collected with priori patient consent signatures and were reviewed and approved by the Ethics Committee of State Key Laboratory of Experimental Hematology, Institute of Hematology and Blood Disease Hospital, Chinese Academy of Medical Sciences and Peking Union Medical College.

## Consent for publication

Not applicable.

## Availability of data and materials

All supporting data are included in the manuscript and supplemental files. Additional data are available upon reasonable request to the corresponding author.

## Authors’ contributions

Y.Q.L. designed and conducted all experiments, performed data analysis and wrote the manuscript. Z.Y.X. performed the core experiments; X.J.Z., C.Y.Z., and F.X.H. performed multiple experiments; Y.W., Y.H.L., H.D.H., Z.Q.W., C.X.X., Q.T.W., L.Q.Z., Y.Q.Z., H.M.Q., Y.Y.S., Y.C., F.Z., J.X.W., P.C.L., J.C.X., L.J.L., and Y.P.Z. assisted in completing the experiments. J.L.Z., W.B.Q., A.B.L., X.F.Z., F.X.H., T.J.W., M.Y.Z., and J.Y.W. discussed the data; T.J.W., M.Y.Z. and J.Y.W. wrote the manuscript; and J.Y.W. designed the project and provided final approval of the manuscript.

## Acknowledgements

We thank members of our team for critical discussion and suggestions.

