## Additional file 1 for "Engineered CRO-CD7 CAR-NK cells derived from pluripotent stem cells avoid fratricide and efficiently suppress human T-cell malignancies"

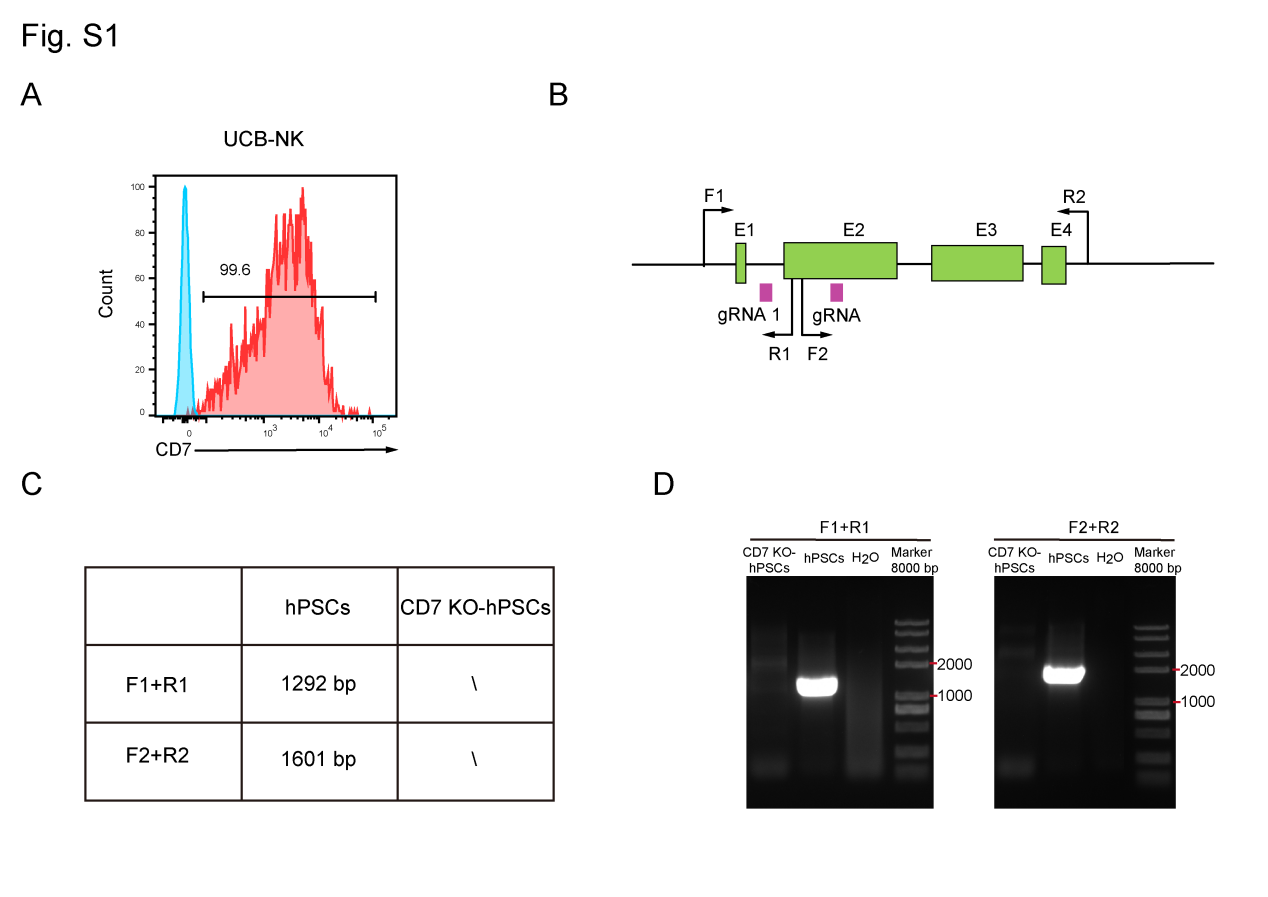

**Fig. S1** **Verification of CD7 knockout in hPSCs. A** Surface expression of CD7 in UCB-NK cells measured by flow cytometry. **B-C** The experimental design of primer pairs to utilize for the identification of CD7 genotypes of hPSCs and CD7 KO-hPSCs. **D** PCR amplification results of each fragment of hPSCs and CD7 KO-hPSCs using the representative primer pairs.

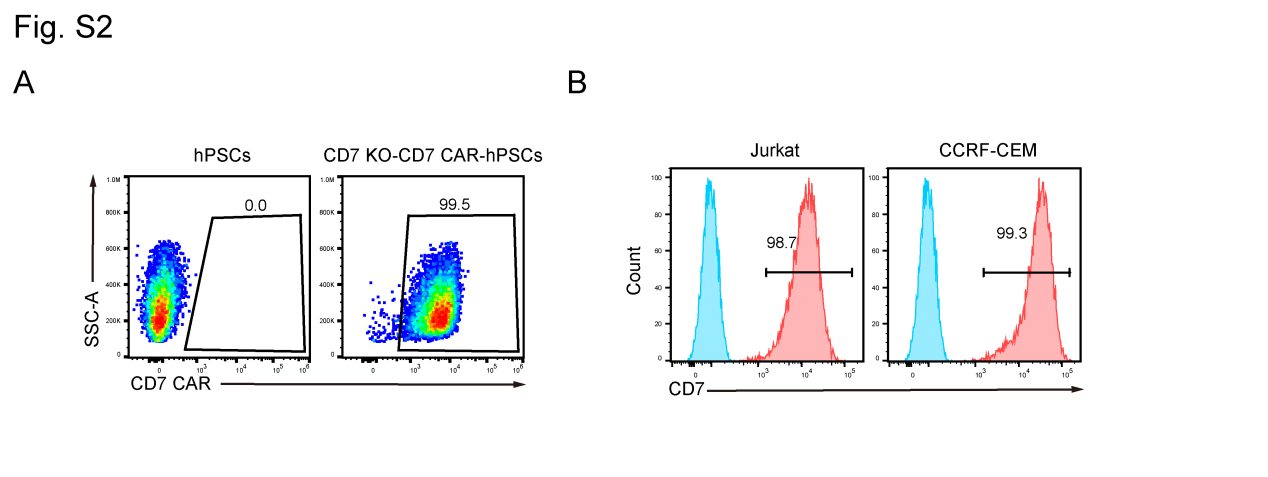

**Fig. S2** **Flow cytometry analysis of the surface expression of CD7 CAR in hPSCs and CD7 KO-CD7 CAR-hPSCs, and of CD7 in Jurkat and CCRF-CEM cells. A** Surface expression of CD7 CAR in hPSCs and CD7 KO-CD7 CAR-hPSCs measured by flow cytometry. **B** Surface expression of CD7 in Jurkat and CCRF-CEM cells measured by flow cytometry.

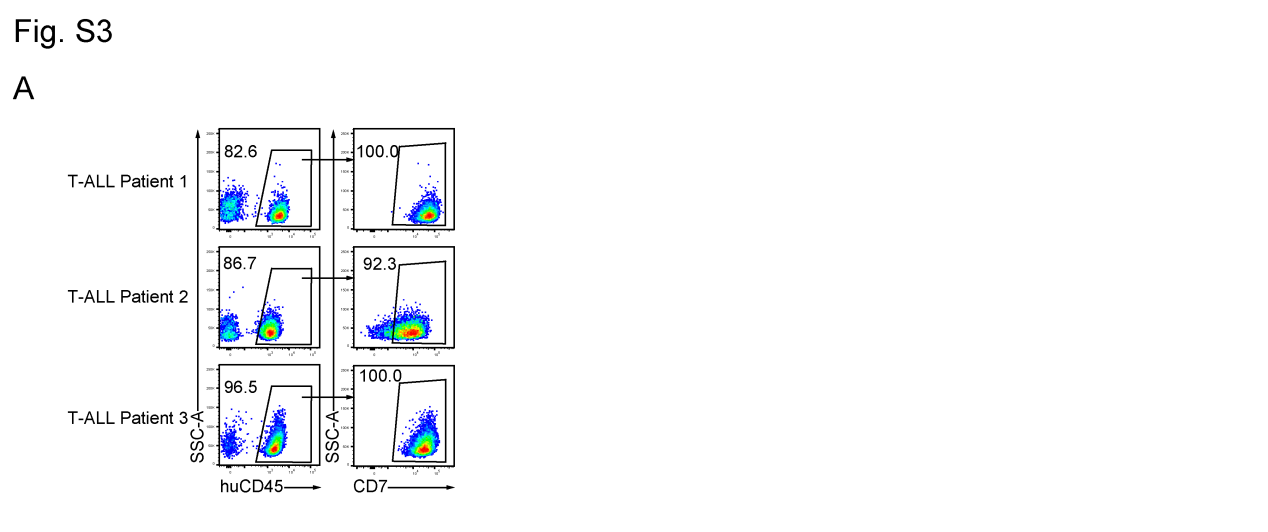

**Fig. S3** **Detection of the proportion of huCD45^+^CD7^+^ cells in the PB of B-NDG mice injected with T-ALL tumor cells. A** Flow cytometry analysis of the huCD45^+^CD7^+^ cells in the PB of B-NDG mice injected with T-ALL tumor cells from patient 1, patient 2, and patient 3, respectively.
